# Direct structural connections between auditory and visual motion selective regions in humans

**DOI:** 10.1101/2020.06.11.145490

**Authors:** A. Gurtubay-Antolin, C. Battal, C. Maffei, M. Rezk, S Mattioni, J. Jovicich, O. Collignon

## Abstract

In humans, the occipital middle-temporal region (hMT+/V5) specializes in the processing of visual motion, while the Planum Temporale (hPT) specializes in auditory motion processing. It has been hypothesized that these regions might communicate directly to achieve fast and optimal exchange of multisensory motion information. In this study, we investigated for the first time in humans the existence of direct white matter connections between visual and auditory motion-selective regions using a combined functional- and diffusion-MRI approach. We found reliable evidence supporting the existence of direct white matter connections between individually and functionally defined hMT+/V5 and hPT. We show that projections between hMT+/V5 and hPT do not overlap with large white matter bundles such as the Inferior Longitudinal Fasciculus (ILF) nor the Inferior Frontal Occipital Fasciculus (IFOF). Moreover, we did not find evidence for the existence of reciprocal projections between the face fusiform area and hPT, supporting the functional specificity of hMT+/V5 – hPT connections. Finally, evidence supporting the existence of hMT+/V5 – hPT connections was corroborated in a large sample of participants (n=114) from the human connectome project. Altogether, this study provides first evidence supporting the existence of direct occipito-temporal projections between hMT+/V5 and hPT which may support the exchange of motion information between functionally specialized auditory and visual regions and that we propose to name the middle (or motion) occipito-temporal track (MOTT).

## INTRODUCTION

Perceiving motion across the senses is arguably one of the most important perceptual skills for the survival of living organisms. Single-cell recordings in primates (1) and functional Magnetic Resonance Imaging (fMRI) studies in humans (2) demonstrated that the middletemporal cortex (hereafter, area hMT+/V5) specializes in the processing of visual motion. One hallmark feature of this region is that it contains cortical columns that are preferentially tuned to a specific direction/axis of visual motion (3). When the function of this region is disrupted, either due to brain damage or by applying transcranial magnetic stimulation, the perception of visual motion directions is selectively impaired (4–6). Even if less research has been dedicated to study the neural substrates of auditory motion, the human planum temporale (hPT), in the superior temporal gyrus, is known to specialize in the processing of moving sounds (7,8). Analogous to hMT+/V5, hPT shows an axis-of-motion organization reminiscent of the organization observed in hMT+/V5 (9). Patients with damage to the right temporal cortex including hPT have shown selective problems in the processing of auditory motion (10,11).

In everyday-life, moving events are often perceived simultaneously across vision and audition. Psychophysical studies have shown the automaticity and perceptual nature of audiovisual motion perception (12–14). In order to create a unified representation of movement, the brain must therefore exchange and integrate motion signals simultaneously captured by vision and audition.

Classical models of movement perception suggest that visual and auditory motion inputs are first processed separately in sensory-specific brain regions and then integrated in multisensory convergence zones (e.g., intraparietal area; (15,16). This hierarchical model has been recently challenged by studies suggesting that the integration of auditory and visual motion information can occur within regions typically considered unisensory (17–20). In particular, in addition to its well-documented role for visual motion, hMT+/V5 has been found to respond preferentially to auditory motion (21) and to contain information about auditory motion directions (22) using a similar representational format in vision and audition (23).

However, how the visual and auditory motion systems communicate is still debated. Although it was initially suggested that audiovisual motion signals in occipital or temporal regions could rely on feedback projections from multimodal areas (24,25), an alternative hypothesis supports the existence of direct connections between motion selective regions in the occipital and temporal cortices (23). This hypothesis finds support in human studies showing increased connectivity between occipital and temporal motion selective areas during the processing of moving information (22,26) as well as in animal tracer studies showing monosynaptic connections between occipital and temporal regions in macaques (27), in particular between motion-sensitive areas (28–31). In humans, the existence of a direct anatomical connections between auditory and visual motion selective regions remains however unexplored.

In our study, we evaluated the presence of direct anatomical connections between hMT+/V5 and hPT using a combined functional and diffusion-weighted MRI approach. To overcome the difficulties in the localization of hMT+/V5 and hPT from anatomical landmarks alone (32,33), each participant was first involved in a visual and an auditory fMRI motion localizer to individually localize these areas functionally. Then, using diffusion MRI data, we reconstructed the connections between these regions by conducting probabilistic tractography and we explored whether connections between hMT+/V5 and hPT followed the trajectories of large white matter bundles such as the Inferior Longitudinal Fasciculus (ILF) or the Inferior Frontal Occipital Fasciculus (IFOF). To further assess the selectivity of hPT – hMT+/V5 connections, we conducted probabilistic tractography between hPT and the Face Fusiform area (FFA), another region of the visual cortex with a specific functional role not related to motion. As an additional control analysis, we investigated the existence of hMT+/V5 – hPT connections in a larger independent dataset (Human Connectome Project) to test the consistency of our findings.

## RESULTS

### Location of individually defined and group-level hMT+/V5 and hPT

Group-level coordinates for hMT+/V5 were located in MNI coordinates (44, −70, −4) and (−50, −70, −2) for right and left hemispheres respectively, which is consistent with reported MNI coordinates for this region (34) and lie within the V5 mask of the Juelich histological atlas available in FSL (35) (see **Figure 1A**). Subject-specific hMT+/V5 coordinates were on average 7 ± 3 (M ± SD) mm and 10 ± 4 mm away from the group-maxima on the right and the left hemisphere (32).

**Figure 1.**
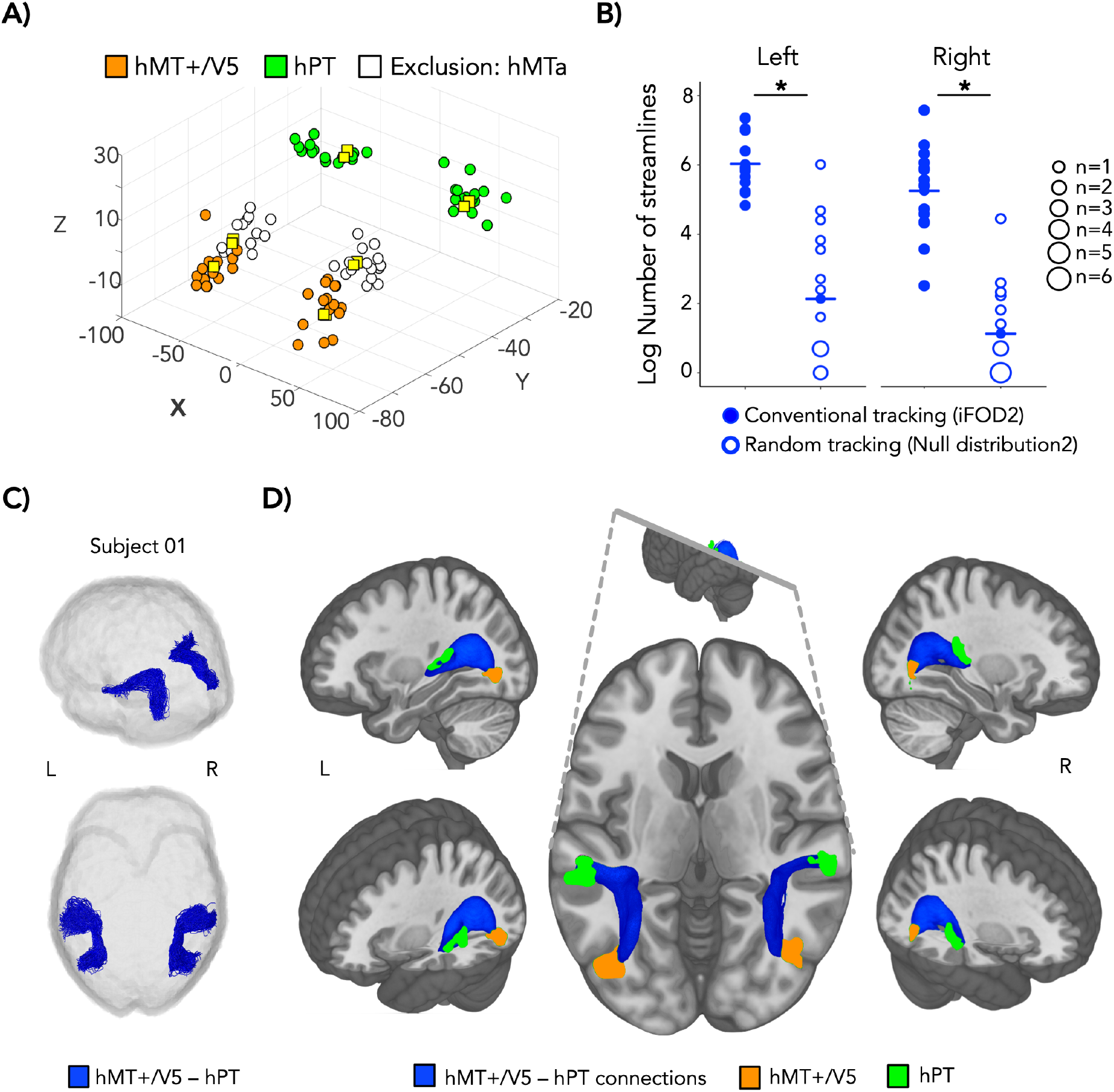
**(A)**. 3D scatterplot depicting subject-specific hMT+/V5 (orange), hPT (green) and exclusion mask (region anterior to hMT+/V5-hMTa-preferentially responding to auditory motion, white) in MNI coordinates. Yellow squares represent the group-level peak-coordinates for the same regions. **(B)**. Balloon plot illustrating the log-transformed number of streamlines reconstructed for hMT+/V5 – hPT connections. Streamlines are reconstructed by conventional tracking (‘iFOD2’ algorithm, represented with filled circles) and random tracking (‘Null distribution2’ algorithm, represented with empty circles). The iFOD algorithm relies on Fiber Orientation Distributions (FODs) whereas the streamlines generated with the Null Distribution algorithm rely on random orientation samples where no information relating to fiber orientations is used. Asterisks represent significant differences. R: right, L: left. **(C).** Resulting tractography reconstruction for hMT+/V5 – hPT connections (blue) for one representative subject. **(D)**. Group-averaged structural pathways between hMT+/V5 and hPT (blue). Inclusion regions hMT+/V5 and hPT are shown in orange and green respectively. Individual connectivity maps were binarized, overlaid and are shown at a threshold of >9 subjects. Inclusion regions followed the same procedure and are shown at a threshold of >2 subjects. Results are depicted on the T1 MNI-152 template. R: right, L: left.

Group-level hPT coordinates were located at coordinates (64, −34, 13) and (−44, −32, 12) for the right and left hemisphere, respectively, which is consistent with reported MNI coordinates for this region (7) and lie within the hPT Harvard–Oxford atlas from FSL (36). Individually defined hPT coordinates were on average 9 ± 5 mm and 15 ± 4 mm away from the group-maxima, for the right and left hemisphere, respectively, which is also consistent with reported inter-individual variability in the location of hMT+/V5 (9).

The respective locations of individually defined coordinates for hMT+/V5, hPT and exclusion region hMTa are illustrated in **Figure 1A**.

### Testing the presence of individually defined hMT+/V5 – hPT connections

For hMT+/V5 – hPT connections that relied on individual hMT+/V5 and hPT, the percentage of streamlines rejected based on aberrant length or position was (M ± SD) 3.9 ± 3.9 for the right and 6.3 ± 4.3 for the left hemisphere. The number of streamlines in all participants was within a range of 3 SDs away from the group mean. For the right hemisphere, the number of reconstructed streamlines was significantly above chance, as the comparison between the connections reconstructed driven by random tracking (‘Null distribution’) with those driven by conventional tracking (‘iFOD2’), revealed a higher streamline count generated with the ‘iFOD2’ algorithm compared to that produced by the ‘Null distribution’ algorithm (log streamlines[iFOD2] = 5.2 ± 1.2, log streamlines[Null distribution] = 1.1 ± 1.3, Paired t-Test, t(14) = 9.7, p = 1e-7, d=2.5). Similar results were obtained for the left hemisphere (log streamlines[iFOD2] = 6.0 ± 0.9, log streamlines [Null distribution] = 2.1 ± 2.0, Paired t-Test, t(14) = 8.3, p = 8e-7, d=2.1). The distribution of the number of streamlines generated with each algorithm can be seen in **Figure 1B**. Tractography reconstruction for hMT+/V5 – hPT connections in a representative subject is illustrated in **Figure 1C** and group-averaged tracts derived are shown in **Figure 1D**.

### Overlap of hMT+/V5 – hPT connections with the IFOF and ILF

The position of hMT+/V5 – hPT connections, as reflected by the sum of binarized individual tract-density images that were thresholded at >9 subjects, relative to the ILF and the IFOF can be seen in **Figure 2A.** The Dice similarity coefficient (DSC) was used as a metric to evaluate the spatial overlap between hMT+/V5 – hPT connections and the Inferior Longitudinal Fasciculus (ILF) and the Inferior Frontal Occipital Fasciculus (IFOF). For the overlap between hMT+/V5 – hPT and the ILF, the DSC was (M ± SD) 0.036 ± 0.027 in the right hemisphere and 0.059 ± 0.045 in the left hemisphere (see **Figure 2B**). For the overlap between hMT+/V5 – hPT and the IFOF, the DSC was (M ± SD) 0.040 ± 0.024 in the right hemisphere and 0.103 ± 0.048 in the left hemisphere. The DSC values were lower when we calculated the overlap between hMT+/V5 – hPT connections, as reflected by the sum of binarized individual tract-density images that were thresholded at >9 subjects, and the ILF (see **Figure 2C**). The DSCs were 7 e-4 for the right and 0.033 for the left hemisphere. The overlap of hMT+/V5 – hPT connections with the IFOF was 8 e-3 for the right and 0.084 for the left hemisphere.

**Figure 2.**
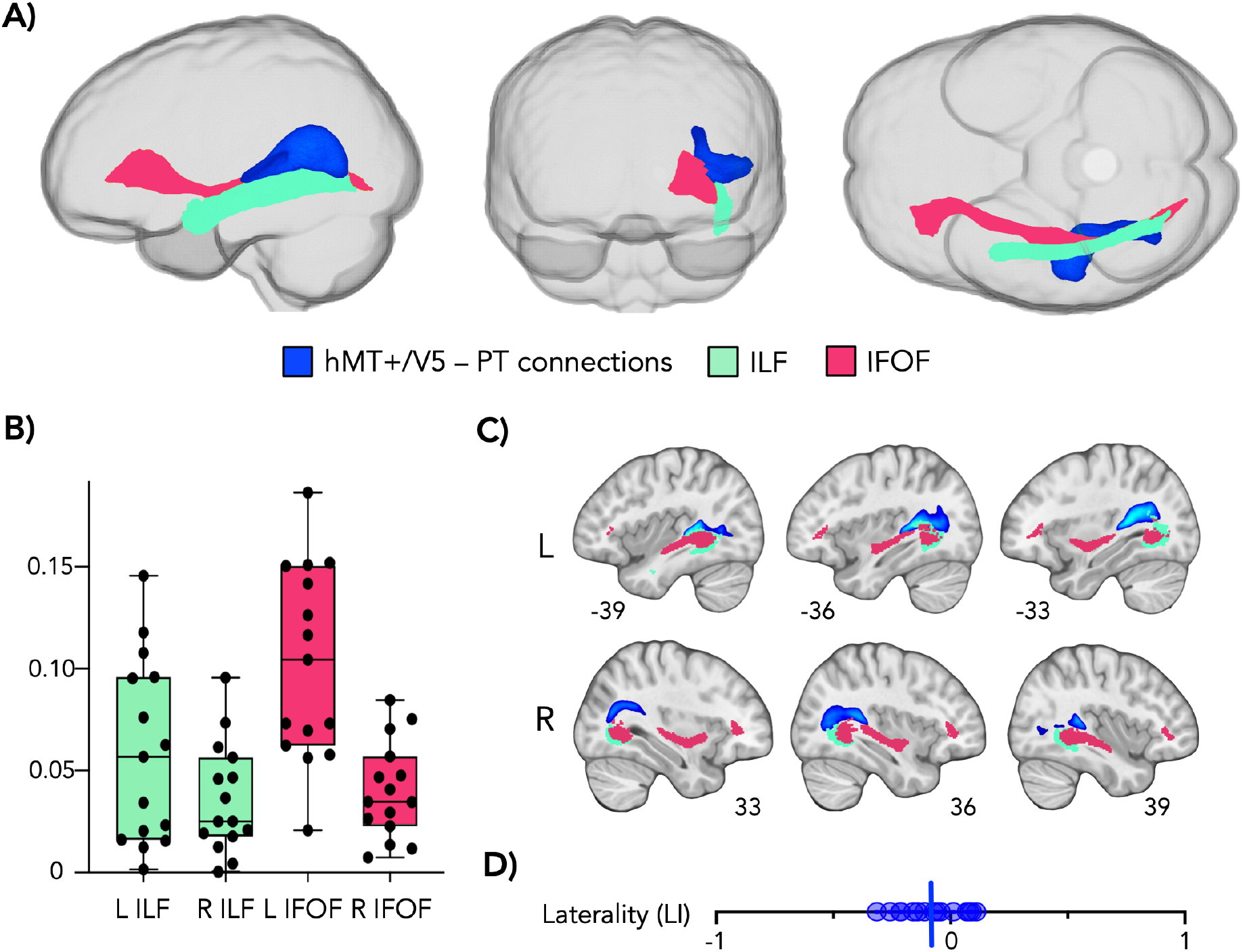
**(A)**. Position of left hMT+/V5 – hPT connections (blue) relative to the left Inferior Frontal Occipital Fasciculus (IFOF, pink) and the left Inferior Longitudinal Fasciculus (ILF, turquoise). Sagittal view (left), coronal view from the anterior part of the brain (middle) and axial view from the inferior part of the brain (right). ILF and IFOF from the JHU-DTI-based white-matter atlas available in FSL have been thresholded at a probability of 25%. hMT+/V5 – hPT connections shown at a threshold of >9 subjects. Results are depicted on the MNI-152 standard space. **(B)** Boxplots represent Dice Similarity Coefficients between individual hMT+/V5 – hPT connections and the ILF (turquoise) and the IFOF (orange). Dots correspond to individual DSC values. **(C)**. Sagittal slices showing the position of hMT+/V5 – hPT connections (threshold of >9 subjects), the ILF and the IFOF in the left (upper row) and right (lower row) hemispheres; R: right, L: left. **(D)**. Laterality index for hMT+/V5 – hPT connections. The laterality index varies between −1 and 1, for completely left- and right-lateralized connections, respectively. Individual values are represented by dots. Vertical line in blue represents the mean value.

### Laterality of hMT+/V5 – hPT connections

hMT+/V5 – hPT connections derived from individually defined hMT+/V5 and hPT were slightly left-lateralized with a the laterality index (M ± SD) −0.082 ± 0.135. The LI in the subject presenting the most left-lateralized connections was −0.315, whereas the LI in the subject presenting the most right-lateralized connections was 0.109. Individual LI values are illustrated in **Figure 2D.**

### Are hMT+/V5 – hPT connections different depending on individual or group-level hMT+/V5 and hPT?

The effect of relying on group-averaged functional data to determine the location of hMT+/V5 and hPT instead of relying on individual activity maps was assessed by three analyses. First, we investigated whether hMT+/V5 – hPT connections existed when the location of the ROIs was derived from group-averaged functional data. We then compared the connectivity index between the tracts derived from group or subject-level functional data. Moreover, microstructural diffusivity measures were contrasted between them.

### Testing the presence of group-level hMT+/V5 – hPT connections

The percentage of streamlines rejected based on aberrant length or position was (M ± SD) 5.0 ± 4.3 for the right and 9.4 ± 2.7 for the left hemisphere. The number of streamlines in all participants was within a range of 3 SDs away from the group mean. hMT+/V5 – hPT connections derived from hMT+/V5 and hPT defined at the group-level were reconstructed above chance suggesting that the dMRI data produced meaningful streamlines between hMT+/V5 and hPT. For right hMT+/V5 – hPT connections, the log-transformed number of streamlines reconstructed with the ‘iFOD2’ algorithm (M ± SD = 6.2 ± 1.3) was significantly higher than those reconstructed by random tracking (M ± SD = 2.3 ± 1.1) (Paired t-Test, t(14) = 11.1, p = 2e-8, d = 2.9). Similar results were obtained for left hMT+/V5 – hPT connections (log streamlines[iFOD2] = 6.2 ± 0.9, log streamlines [Null distribution] = 4.4 ± 0.8, Paired t-Test, t(14) = 6.9, p = 8e-6, d = 1.8). The overlap between hMT+/V5 – hPT connections derived from group-level or subject-level functional data is shown in **Figure 3A**.

**Figure 3.**
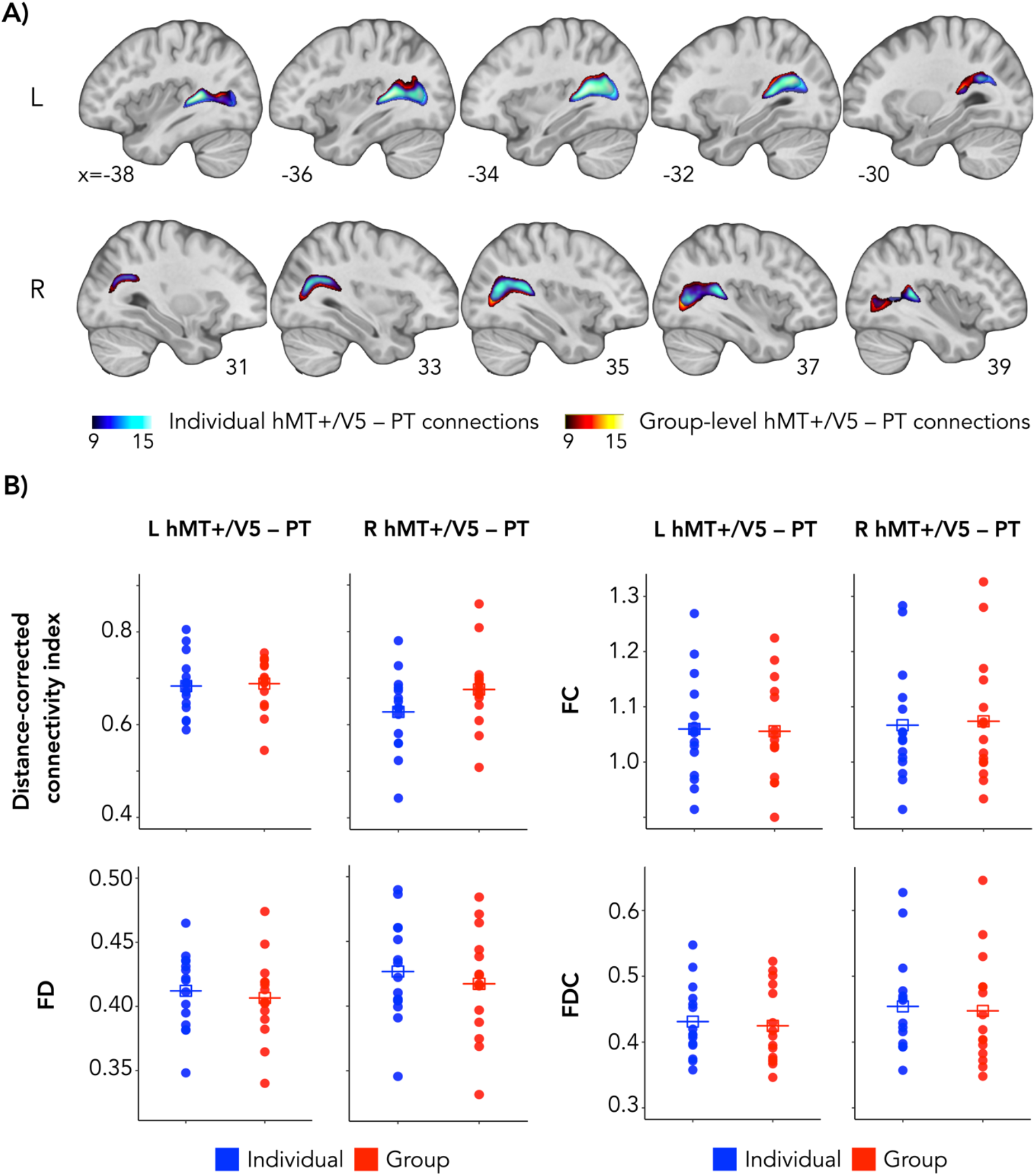
**(A)** Overlap of group-averaged structural pathways between hMT+/V5 and hPT for connections derived from individual ROIs (blue) and group-level ROIs (red). Squares crossed by horizontal lines represent mean values. Individual connectivity maps were binarized and overlaid. Bar-color codes the number of subjects showing the path in each voxel (thresholded at >9 subjects). Tracts derived from individual ROIs (blue) are shown at 75% opacity for better visualization of the overlap. Results are depicted on the T1 MNI-152 template. R: right, L: left. **(B)** Distance-corrected connectivity index and microstructural metrics FD, FC and FDC, for right and left hMT+/V5 – hPT connections derived from individual (Individual, blue) or group-level (Group, red) hMT+/V5 and hPT. The connectivity index has been corrected for distance. R: right, L: left.

### Connectivity index

For right hMT+/V5 – hPT connections, the connectivity index (CI) for tracts derived from individual ROIs (M ± SD = 0.37 ± 0.09) was lower than that obtained from group-level ROIs (M ± SD = 0.42 ± 0.09) (Paired t-Test, t(14) = 2.77, p = 0.01, d = 0.7). For left hMT+/V5 – hPT connections, the CI between tracts relying on subject-specific or group-level ROIs revealed no differences (CI [subject-specific ROIs] = 0.42 ± 0.06, CI [group-level ROIs] = 0.44 ± 0.06, Paired t-Test, t(14) = 0.87, p = 0.4, d = 0.2).

To control for possible differences in the distance between seed and targets that could drive differences in the connectivity index, we compared the seed-target distance between individual or group-level ROIs. For both right and left hMT+/V5 – hPT connections, the distance between hMT+/V5 and hPT was larger when relying on individually defined ROIs than for group-level ROIs (right hemisphere: log Distance [subject-specific ROIs] = 3.71 ± 0.11, log Distance [group-level ROIs] = 3.60 ± 0.08, Paired t-Test, t(14) = −2.93, p = 0.01, d = 0.8; left hemisphere: log Distance [subject-specific ROIs] = 3.71 ± 0.20, log Distance [group-level ROIs] = 3.57 ± 0.07, Paired t-Test, t(14) = −2.81, p = 0.01, d = 0.7). Hence, we computed the distance-corrected connectivity index, replacing the number of streamlines, by the product of the number of streamlines and the distance between the hMT+/V5 and hPT. Distance-corrected connectivity indexes did not differ between individual or group-level ROIs for any hemisphere (right hemisphere: CI [subject-specific ROIs] = 0.63 ± 0.08, CI [group-level ROIs] = 0.68 ± 0.08, Paired t-Test, t(14) = 2.5, p = 0.03, d = 0.6; left hemisphere: CI [subject-specific ROIs] = 0.68 ± 0.06, CI [group-level ROIs] = 0.68 ± 0.06, Paired t-Test, t(14) = 0.28, p = 0.8, d =0.07). This means that the lower connectivity index observed for individual ROIs compared to group-level ROIs in the right hemisphere, was likely due to a higher distance between individual hMT+/V5 and hPT. The distribution of the distance-corrected connectivity index for tracts derived from subject-specific or group-level hMT+/V5 and hPT can be seen in **Figure 3B**.

### FOD-derived microstructural metrics

Tissue microstructure was addressed by within-voxel fiber density (FD), fiber-bundle crosssection (FC) and their combined measure (FDC). Fiber density of streamlines connecting hMT+/V5 and hPT did not differ when relying on individual ROIs compared to group-level ROIs, in any hemisphere (right hemisphere: FD [subject-specific ROIs] = 0.43 ± 0.04, FD [group-level ROIs] = 0.42 ± 0.04, Paired t-Test, t(14) = −1.89, p = 0.08, d = 0.5; left hemisphere: FD [subject-specific ROIs] = 0.41 ± 0.09, FD [group-level ROIs] = 0.41 ± 0.09, Paired t-Test, t(14) = −1.06, p = 0.3, d = 0.3). See **Figure 3B** for the distribution of microstructural diffusivity metrics for tracts derived from subject-specific or group-level hMT+/V5 and hPT. The fiber-bundle cross-sectional area (FC) did not differ between tracts derived from individual or group-level functional data, suggesting an absence of difference in the number of voxels that the fiber-bundle occupies (right hemisphere: FC [subject-specific ROIs] = 1.07 ± 0.11, FC [group-level ROIs] = 1.07 ± 0.11, Paired t-Test, t(14) = 1.27, p = 0.2, d = 0.3; left hemisphere: FC [subject-specific ROIs] = 1.06 ± 0.10, FC [group-level ROIs] = 1.06 ± 0.10, Paired t-Test, t(14) = −0.95, p = 0.4, d = 0.2). Finally, fiber density and cross-section (FDC), which describes changes in the total intra-axonal volume by combining the previous two metrics, was neither found to differ between tracts that relied on subject-specific or group-level ROIs (right hemisphere: FDC [subject-specific ROIs] = 0.45 ± 0.08, FDC [group-level ROIs] = 0.44 ± 0.08, Paired t-Test, t(14) = −0.99, p = 0.3, d = 0.3; left hemisphere: FDC [subject-specific ROIs] = 0.43 ± 0.05, FDC [group-level ROIs] = 0.42 ± 0.06, Paired t-Test, t(14) = −0.97, p = 0.3, d = 0.2).

### Testing the presence of group-level FFA – hPT connections

The percentage of streamlines discarded based on the length or position criteria was (M ± SD) 9.2 ± 2.6 for the right and 10.2 ± 3.2 for the left hemisphere. All the participants were included in the analysis, as there were no outliers considering the number of streamlines. As opposed to hMT+/V5 – hPT connections, we did not find evidence to suggest the existence of FFA – hPT connections. For the right hemisphere, the number of streamlines derived from a null distribution did not differ compared to the number of streamlines generated with the ‘iFOD’ algorithm (right FFA – hPT (log streamlines[iFOD2] = 5.0 ± 1.5, log streamlines [Null distribution] = 5.1 ± 1.1, Paired t-Test, t(14) = −0.6, p = 0.6, d = 0.2). The absence of difference between the number of streamlines derived from a null distribution and the number of streamlines generated with the ‘iFOD’ algorithm, was further assessed by the Bayes factor (BF). Low BF values (<1) represent how strongly the data supports the null hypothesis of no effect. In accordance with the previous results, the BF was 0.307.

For the left hemisphere, the number of streamlines derived from a null distribution was higher than the number of streamlines obtained with the ‘iFOD’ algorithm, showing an opposite pattern compared to the results obtained for hMT+/V5 – hPT connections (log streamlines[iFOD2] = 5.0 ± 1.2, log streamlines [Null distribution] = 6.4 ± 1.0, Paired t-Test, t (14) = −6.9, p = 7e-6, d = 1.8). This means that diffusion data is not providing evidence for the existence of FFA-hPT connections than that expected from random tracking.

### Replication in Dataset 2: Testing the presence of hMT+/V5 – hPT connections

To evaluate the consistency of hMT+/V5 – hPT connections obtained in the Dataset 1, the existence of reciprocal hMT+/V5 – hPT projections was also investigated in the Human Connectome Project dataset (Dataset 2). This dataset allowed us to test the reproducibility of our findings using a larger sample size (as compared to Dataset 1) and different acquisition parameters. Whereas single-shell diffusion data was acquired in Dataset 1, multi-shell diffusion data was acquired in Dataset 2. For the reconstruction of hMT+/V5 – hPT tracts on this dataset we relied on the group-level hMT+/V5 and hPT described in the Dataset 1. We corroborated the existence of hMT+/V5 – hPT connections above chance levels. For right hMT+/V5 – hPT connections, the percentage of streamlines rejected based on aberrant length or position was (M ± SD) 3.4 ± 4.2. The number of streamlines (log-transformed) were not normally distributed (Shapiro-Wilk test, p=1e-7). Hence, in addition to a paired t-Test we also assessed differences using the non-parametric Wilcoxon test. The log-transformed number of streamlines reconstructed with the ‘iFOD2’ algorithm (M ± SD = 1.92 ± 1.86) was significantly above chance (M ± SD = 0.04 ± 0.17) after removing three outlier participants whose number of streamlines was more than 3 SD away from the group mean (Paired t-Test, t(110) = 10.9, p = 2.2e-16, d = 1.0; Wilcoxon Test, Z = −7.47, p = 8e-14, n=111). For left hMT+/V5 – hPT connections, the percentage of streamlines discarded based on aberrant length or position was (M ± SD) 5.2 ± 4.9. The log-transformed number of streamlines were normally distributed and streamlines were reconstructed above chance-levels after rejecting one participant due to aberrant number of streamlines (log streamlines[iFOD2] = 2.94 ± 2.27, log streamlines [Null distribution] = 2.16 ± 1.64, paired t-Test, t(113) = 4.4, p = 2e-5, d = 0.4). Group-averaged tracts derived for this dataset can be seen in **Figure 4A** and the overlap between hMT+/V5 – hPT connections from Dataset 1 and Dataset 2 is shown in **Figure 4B**.

**Figure 4.**
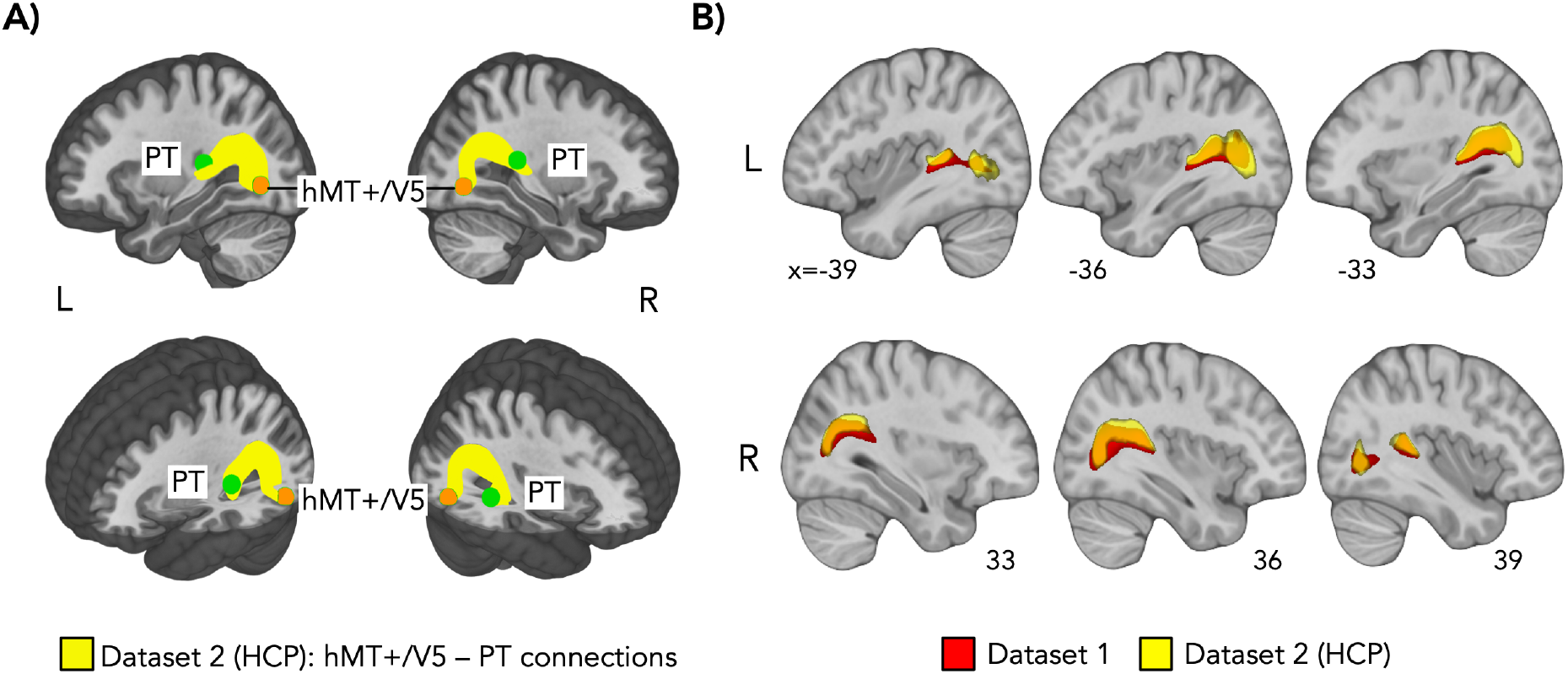
**(A)** Group-averaged structural pathways between hMT+/V5 and hPT (yellow) in Dataset 2 (HCP). Inclusion regions hMT+/V5 and hPT are shown in orange and green respectively. Individual connectivity maps were binarized, overlaid and are shown at a threshold of >9 subjects. Results are depicted on the T1 MNI-152 template. R: right, L: left. **(B)** Overlap of hMT+/V5 – hPT connections (from group-level ROIs) derived from Dataset 1 (red) and Dataset 2 (yellow). Tracts derived from Dataset 2 are shown at 75% opacity for better visualization of the overlap. Results are depicted on the T1 MNI-152 template. R: right, L: left.

## DISCUSSION

We investigated for the first time the potential existence of direct white matter connections in humans between visual and auditory motion-selective regions using a combined functional- and diffusion-MRI approach. Diffusion imaging can be used to estimate brain connectivity in-vivo (37), which can be analyzed alongside fMRI data for the same individual and therefore relate structure and function. We found reliable evidence for the existence of a direct pathways between functionally defined hMT+/V5 and hPT in two independent datasets. We show that the projections between these regions do not follow the trajectories of other large white matter bundles such as the ILF or the IFOF. In contrast, we did not find evidence to support the existence of reciprocal projections between the fusiform face area and hPT, suggesting that direct connections only emerge between temporal and occipital regions that share a similar computational goal. Our findings suggest the existence of direct occipito-temporal pathways between hMT+/V5 and hPT in humans that might support a fast and optimal integration of auditory and visual motion information (26).

Visual and auditory motion signals interact to create a unified perception of motion (25). Stimuli moving in opposite directions across the senses lead to erroneous motion perception (13), whereas congruent audio-visual moving stimuli are shown to enhance the processing of directional motion (19,26,38,39). Moreover, perceiving the direction of visual motion can lead an ambiguous auditory stimulus to be perceived as moving in the opposite direction (crossmodal adaptation) (12). But which is the neural architecture supporting the integration of multisensory motion information? Classical hierarchical models of information processing in the brain suggest that visual and auditory motion inputs are first processed separately in sensory-selective regions and then integrated in multisensory convergence zones like the posterior parietal and premotor cortices (15). In contrast to this purely hierarchical view of how multisensory motion information unfold in the brain, moving sounds have been found to modulate the spike-count rate of neurons responding to visual motion (40) and influence the BOLD response of hMT+/V5 (19,26,38,41). In fact, moving sounds activate the anterior portion of MT+/V5 in humans and macaques (21,23,42) and planes of auditory motion are successfully encoded in the distributed activity of hMT+/V5 (22). Importantly, motion directions in the visual modality can be predicted by the patterns elicited by audition directions in hMT+/V5, and reversely, demonstrating a partially shared representation for motion direction across the senses in hMT+/V5 (23).

Initially, it was proposed that the presence of auditory motion signal in hMT+/V5 results from feed-back connection from higher-order convergence zones like the posterior parietal cortex (16). However, the exclusive role of feedback connections in the integration of audiovisual motion in regions typically conceived as unisensory has been challenged by studies showing audiovisual interactions as early as 40 ms after stimulus presentation (18,43,44). Moreover, tracer studies in animals have demonstrated the existence of direct monosynaptic connections between primary auditory and visual regions (27,47). In agreement with our findings, previous macaque studies have revealed monosynaptic connections between area MT, the equivalent of area hMT+/V5 in primates, and regions in the temporal lobe assumed to be sensitive to auditory motion (28–31). A recent study further revealed the existence of direct projections from the caudal portions of the middle lateral belt region of the auditory cortex, thought to represent part of the hPT equivalent in macaques (42), to area MT (48). Our suggestion for direct reciprocal projections between hMT+/V5 and hPT is consistent with those findings highlighting the existence of cortical pathways for sensory information exchange at a much lower level of the information processing hierarchy than was previously thought (45,46,49).

We investigated the specificity of the hMT+/V5 – hPT pathway. First, we showed that hMT+/V5 – hPT connections do not follow the same trajectory as neither the inferior longitudinal fasciculus (ILF), the major occipito-temporal white matter bundle (50,51), nor the inferior frontal occipital fasciculus (IFOF), the main direct connection between the occipital and frontal lobes. To further investigate the selectivity of the hMT+/V5 – hPT projections, we investigated the existence of projections between hPT and the FFA, another functionally selective region in the occipital cortex that preferentially responds to faces (52). Interestingly, previous studies have shown direct connections between FFA and a region in the middle temporal gyrus selective to vocal sounds (53,54). In accordance with a specific role of hMT+/V5 – hPT connections in motion processing, we did not find evidence to suggest the existence of connections between FFA and hPT. Altogether, our results suggest that direct connections only emerge between temporal and occipital regions that share a similar computational goal (e.g. in our study regarding moving information) since these regions need a fast and optimal transfer of redundant perceptual information across the senses. The intrinsic preferential connectivity between functionally related brain regions such as hMT+/V5 and hPT for motion could provide the structural scaffolding for the subsequent development of these areas in a process of interactive specialization (55). Evolution may have coded in our genome a pattern of connectivity between functionally related regions across the senses, as for instance those involved in motion perception, to facilitate the emergence of functional networks dedicated to a perceptual/cognitive process and facilitate exchange in computationally related multisensory information. Actually developmental studies show that young infants (56) and even newborns (57) can detect motion congruency across the senses.

Interestingly, these intrinsic connections between sensory systems devoted to the processing of moving information may constrain the expression of crossmodal plasticity in case of early sensory deprivation. Several studies have indeed reported that early blinds preferentially recruit an area consistent with the location of hMT+/V5 for the processing of moving sounds (22,58,59). Likewise, hPT region typically sensitive to moving sounds, is preferentially recruited by visual motion in case of early deafness (60–62). In this context, crossmodal reorganization would express relying on occipito-temporal motion selective connections that can also be found in people without sensory deprivation (63,64), such as the hMT+/V5 – hPT connections that we find in the present study.

Despite the relevance of diffusion MRI to support the presence of white matter bundles that are reconstructed by following pathways of high diffusivity (65), the reconstruction of streamlines from diffusion MRI is however an imperfect attempt to capture the full complexity of millions of densely packed axons that form a bundle (66,67). As such, diffusion imaging in humans cannot be considered as a definitive proof for the existence of an anatomical pathway in the brain (68). As for all studies involving diffusion MRI data, postmortem tracer studies could help to overcome the inherent constrains of diffusion MRI, providing direct evidence for the existence of a white matter connection as well as information on the directionality of the connections and finer spatial details (69–71).

In summary, our findings suggest the existence of a direct occipito-temporal pathway connecting hMT+/V5 and hPT, that could represent the structural scaffolding for the exchange and integration of visual and auditory motion information. We propose to name this putative pathway the middle (or motion) occipito-temporal track (MOTT). This finding has important implications for our understanding on how multisensory information is shared across the senses, pointing to the existence of computationally specific pathways that allow information flow between areas traditionally conceived as unisensory, in addition to the bottom-up integration of sensory signals in higher-order multisensory areas.

## MATERIALS AND METHODS

### Dataset 1: Trento

16 participants (6 women; mean age ± SD, 30.6 ± 5.1; range, 20–40) were scanned at the Center for Mind/Brain Sciences (CIMeC) of the University of Trento using a Bruker BioSpin MedSpec 4T MR-scanner equipped with 8-channel transmit and receive head coil. The study was approved by the Committee for Research Ethics of the University of Trento. All participants gave informed consent in agreement with the ethical principles for medical research involving human subject (Declaration of Helsinki, World Medical Association) and the Italian Law on individual privacy (D.l. 196/2003). One participant was excluded from the analysis due to excessive movement during the auditory motion localizer task.

#### Image acquisition

Four imaging datasets were acquired from each participant: a functional MRI visual motion localizer, a functional MRI auditory motion localizer, diffusion-weighted MR images and structural T1-weighted images. Participants were instructed to lie still during acquisition, and foam padding was used to minimize scanner noise and head movement.

##### Functional MRI (fMRI) sequences

Functional images were acquired with T2*-weighted gradient echoplanar sequence. Acquisition parameters were as follows: repetition time (TR), 2500 ms; echo time (TE), 26 ms; flip angle (FA), 73°; field of view, 192 mm; matrix size, 64 x 64; and voxel size, 3 x 3 x 3 mm^3^. A total of 39 slices was acquired in an ascending feet-to-head interleaved order with no slice gap. The three initial scans of each acquisition run were discarded to allow for steady-state magnetization. Before each EPI run, we performed an additional scan to measure the pointspread function (PSF) of the acquired sequence, including fat saturation, which served for distortion correction that is expected with high-field imaging (72).

###### Functional visual motion localizer experiment

A visual motion localizer experiment was implemented to localize hMT+/V5. Visual stimuli were back-projected onto a screen (width: 42 cm, frame rate: 60 Hz, screen resolution: 1024 x 768 pixels; mean luminance: 109 cd/m^2^ via a liquid crystal projector (OC EMP 7900, Epson Nagano) positioned at the back of the scanner and viewed via mirror mounted on the head coil at a distance of 134 cm. Stimuli were 16 s of random-dot patterns, consisting of circular aperture (radius 4°) of radial moving and static dots (moving and static conditions, respectively) with a central fixation cross (73). One hundred and twenty white dots (diameter of each dot was 0.1 visual degree) were displayed on a gray background, moving 4° per second. In all conditions, each dot had a limited lifetime of 0.2 s. Limited lifetime dots were used in order to ensure that the global direction of motion could only be determined by integrating local signals over a larger summation field rather than by following a single dot (74). Additionally, limited lifetime dots allowed the use of control flickering (as opposed to purely static). Stimuli were presented for 16 s followed by a 6 s rest period. Stimuli within motion blocks alternated between inward and outward motion (expanding and contracting) once per second. Because the localizer aimed to localize the global hMT+/V5 complex (e.g. MT and MST regions) the static block was composed of dots maintaining their position throughout the block in order to prevent flicker-like motion (75). The localizer consisted of 14 alternating blocks of moving and static dots (7 each) and lasting a total of 6 m 40 s (160 volumes). In order to maintain the participant’s attention and to minimize eye-movement during acquisition during the localizer’s run, participants were instructed to detect a color change (from black to red) of a central fixation cross (0.03°) by pressing the response button with the right index finger. Three out of the sixteen participants had a slightly modified version of such a visual motion localizer as described elsewhere (23).

###### Functional auditory motion localizer experiment

To localize hPT, we implemented an auditory motion localizer. To create an externalized ecological sensation of sound location and motion inside the MRI scanner, we recorded individual in-ear stereo recordings in a semi-anechoic room outside the MRI scanner and on 30 loudspeakers on horizontal and vertical planes, mounted on two semicircular wooden structures. Participants were seated in the center of the apparatus with their head on a chinrest, such that the speakers on the horizontal and vertical planes were equally distant from participants’ ears. Sound stimuli consisted of 1250 ms pink noise (50 ms rise/fall time). In the motion condition, pink noise was presented moving in leftward, rightward, upward, and downward directions. In the static condition, the same pink noise was presented separately at one of the following four locations: left, right, up, and down. These recordings were then replayed to the blindfolded participants inside the MRI scanner via MR-compatible closed-ear head-phones (500–10 KHz frequency response; Serene Sound, Resonance Technology). In each run, participants were presented with eight auditory categories (four motion and four static) randomly presented using a block design. Each category of sound was presented for 15 s [12 repetitions of 1250 ms sound, no interstimulus interval (ISI)] and followed by 7 s gap to indicate the corresponding direction/location in space and 8 s of silence (total interblock interval, 15 s). Participants completed a total of 12 runs, with each run lasting 4 min and 10 s. A more detailed description can be found elsewhere (9). As for the visual modality, three participants conducted a slightly modified version of this auditory motion localizer that had one long run of 13 motion blocks and 13 static blocks (587.5 secs in total). For more details, see (23).

##### Diffusion MRI (dMRI)

Whole brain diffusion weighted images were acquired using an EPI sequence (TR = 7100 ms, TE = 99 ms, image matrix = 112 × 112, FOV = 100 × 100 mm^2^, voxel size 2.29 mm isotropic). Ten volumes without any diffusion weighting (b0-images) and 60 diffusion-weighted volumes with a b-value of 1500 s/mm2 were acquired. By using a large number of directions and ten repetitions of the baseline images on a high magnetic field strength we aimed at improving the signal-to-noise ratio and reduce implausible tracking results (76).

##### Structural (T1) images

To provide detailed anatomy a total of 176 axial slices were acquired with a T1-weighted MP-Rage sequence covering the whole brain. The imaging parameters were: TR = 2700 ms, TE= 4.18ms, flip angle = 7°, isotropic voxel = 1 mm3, field of view, 256 x 224 mm and inversion time, 1020 ms (77).

#### Image processing

##### Definition of regions of interest (ROIs) using functional data

Functional volumes from the localizer experiments were used to define regions responding preferentially to motion (hMT+/V5 for vision; hPT for audition). For the preprocessing and analysis of functional data we used SPM8 (Welcome Department of Imaging Neuroscience, London), implemented in MATLAB 2016b (MathWorks). The preprocessing of functional data included the realignment of functional time series with a 2nd degree B-spline interpolation, co-registration of functional and anatomical data and spatial smoothing (Gaussian kernel, 6 mm full-width at half-maximum, FWHM). Visual and auditory motion localizer experiments were analyzed separately and the ROIs in each experiment were localized both (1) in each subject individually and (2) at the group level. The rationale behind defining group-level ROIs was twofold: 1) it allowed us to assess the reproducibility of the connection using subject-specific vs. group-level ROIs and 2) it allowed us was to use the group coordinates in other datasets where the individual localization of hMT+/V5 or hPT is not possible, such as the HCP dataset (Dataset 2; see below).

###### Individually defined hMT+/V5

Following the preprocessing steps, we obtained blood oxygenation level-dependent activity related to visual motion and visual static blocks. For each subject, we computed statistical comparisons with a fixed-effect general linear model (GLM). The GLM was fitted for every voxel with the visual motion and the visual static conditions as regressors of interest and six head motion parameters derived from realignment of the functional volumes (three translational motion and three rotational motion parameters) as regressors of no interest. Each regressor was modeled with a boxcar function and convolved with the canonical hemodynamic response function (HRF). A high-pass filter of 128 s was used to remove low frequency signal drifts. Brain regions that responded preferentially to the moving visual stimuli were identified for every subject individually by subtracting visual motion conditions [Visual Motion] and static conditions [Visual Static]. Subject-specific hMT+/V5 coordinates were defined by identifying voxels in a region nearby the intersection of the ascending limb of the inferior temporal sulcus and the lateral occipital sulcus (32) that responded significantly more to motion than static conditions using a lenient threshold of p<0.01 uncorrected in order to localize this peak in every participant.

###### Group-level hMT+/V5

The preprocessing of the functional data for the group level analysis additionally included the spatial normalization of anatomical and functional data to the to the Montreal Neurological Institute (MNI) template using a resampling of the structural and functional data to an isotropic 2 mm resolution. The individual [Visual Motion > Visual Static] contrast was further smoothed with a 8-mm FWHM kernel and entered into a random effects model for the second-level analysis consisting of a 1 sample t-test against 0. Group-level hMT+/V5 coordinate was defined by identifying voxels in a region nearby the intersection of the ascending limb of the inferior temporal sulcus and the lateral occipital sulcus (32) that responded significantly more to motion than static conditions using familywise error (FWE) correction for multiple comparisons at p<0.05.

###### Individually defined hPT

For the auditory motion localizer, the same preprocessing as for the visual motion localizer was applied. The GLM included eight regressors from the motion and static conditions (four motion directions, four sound source locations) and six movement parameters (three translational motion and three rotational motion parameters) as regressors of no interest. Each regressor was modeled with a boxcar function, convolved with the canonical HRF and filtered with a high-pass filter of 128 s. Brain regions responding preferentially to the moving sounds were identified for every subject individually by subtracting all motion conditions [Auditory Motion] and all static conditions [Auditory Static]. Individual hPT coordinates were defined at the peaks in the superior temporal gyrus that lie posterior to Hesclh’s gyrus and responded significantly more to motion than static. We used a lenient threshold of p<0.01 uncorrected in the individual [Auditory Motion > Auditory Static] contrast to be able to localize this region in each participant.

###### Group-level hPT

For hPT defined at the group-level, an analogous procedure to the one conducted for the visual motion localizer was used.

###### FFA coordinate definition from previous literature

To define the FFA, we relied on face-preferential group coordinates extracted from a previous study of our laboratory (78) for the right [44, −50, −16] and the left [−42, −52, −20] hemisphere (in MNI space).

##### dMRI preprocessing

Data preprocessing was implemented in MRtrix 3.0 (79) (www.mrtrix.org), and in FSL 5.0.9 (http://fsl.fmrib.ox.ac.uk/fssl/fslwiki). Briefly, data were denoised (80), removed of Gibbs-ringing, corrected for Eddy currents distortions and head motion (81) and for low-frequency B1 field inhomogeneities (82). Spatial resolution was up-sampled by a factor of 2 in all three dimensions using cubic b-spline interpolation, to a voxel size of 1.15 mm^3^ and intensity normalization across subjects was performed by deriving scale factors from the median intensity in select voxels of white matter, grey matter, and CSF in b = 0 s/mm^2^ images, then applying these across each subject image (83). This step normalizes the median white matter b = 0 intensity (i.e. non-diffusion-weighted image) across participants so that the proportion of one tissue type within a voxel does not influence the diffusion-weighted signal in another. The T1-weighted structural images were non-linearly registered to the diffusion data in ANTs (84) using an up-sampled FA map (1×1×1 mm3) and segmented in maps for white matter, grey matter, CSF, and sub-cortical nuclei using the FAST algorithm in FSL (85). This information was combined to form a five tissue type image to be used for anatomically constrained tractography in MRtrix3 (86). These maps were used to estimate tissue-specific response functions (i.e. the signal expected for a voxel containing a single, coherently-oriented fiber bundle) for grey matter, white matter, and CSF using Multi-Shell Multi-Tissue Constrained Spherical Deconvolution (MSMT) (CSD) (87). Fiber orientation distribution functions (fODFs) were then estimated using the obtained response function coefficients averaged across subjects to ensure subsequent differences in fODFs amplitude will only reflect differences in the diffusion-weighted signal. Note that by using MSMT-CSD in our single-shell, data benefitted from the hard non-negativity constraint, which has been observed to lead to more robust outcomes (87). Spatial correspondence between participants was achieved by generating a group-specific population template with an iterative registration and averaging approach using FOD images from all the participants (88). This is, each subject’s FOD image was registered to the template using FOD-guided nonlinear registrations available in MRtrix (89). These registration matrices were required to transform the seed and target regions from native diffusion space to the template diffusion space, where tractography was conducted. We chose to conduct tractography in the template diffusion space, as FOD-derived metrics of microstructural diffusivity can only be computed in that space. Subsequently, we extracted the following quantitative measures of microstructural diffusivity for all the fixels (fiber populations within a voxel) in the brain: fiber density (FD), fiber-bundle cross-section (FC) and a combined measure of fiber density and cross-section (FDC) (90). For further details on these metrics, see “FOD-derived microstructural metrics” section.

##### Preparation of ROIs for tractography

###### Individually defined hMT+/V5 and hPT

We transformed individually-defined coordinates from the native structural space to the native diffusion space. The reconstruction of white matter connections between functionally defined regions is particularly challenging because it is likely to encompass portions of grey matter which suffer from ill-defined fiber orientations (91,92). Therefore, once in native diffusion space, the peak-coordinates were moved to the closest white matter voxel (FA>0.25) (53,54) and a sphere of 5 mm radius was centered there. To ensure that tracking was done from white matter voxels only, we masked the sphere with individual white matter masks. Last, ROIs were transformed from native diffusion space to template diffusion space, where tractography was conducted.

###### Group-level hMT+/V5, hPT and FFA

First, we computed the warping images between the standard MNI space and the native structural space of each participant by conducting a non-linear registration in ANTs, between each subject’s T1-images and the MNI152 standard-space T1-template-image. Using these registration matrices, we transformed the group peak-coordinates from the standard MNI space to the native structural space of each participant. Once the coordinates where in native structural space, we followed the same procedure described for the individually defined coordinates.

##### Probabilistic tractography

Probabilistic tractography was conducted between three pairs of ROIs: 1) individually defined hMT+/V5 and hPT, 2) group-level hMT+/V5 and hPT and 3) group-level hPT and FFA. We selected hMT+/V5 and hPT as inclusion regions for tractography, based on their preferential response to visual and auditory motion respectively. In addition to the hPT, a region just anterior to hMT+/V5 (called hMTa; see (23)) is also selectively recruited for the processing of moving sounds (9,21,23,93). To prevent hPT from connecting with regions in the vicinity of hMT+/V5 which respond preferentially to auditory-but not visual-motion, this region anterior to hMT+/V5 that respond selectively to moving sound (hMTa; identified in the auditory motion localizer task) were used as an exclusion mask for connections between hMT+/V5 and hPT. This way, we avoided the reconstruction of tracts between motion selective regions responding preferably to the auditory modality.

For each pair of ROIs, we computed tractography in symmetric mode (i.e. seeding from one ROI and targeting the other, and conversely). We then merged the tractography results pooling together the reconstructed streamlines. We used two tracking algorithms in MRtrix (‘iFOD2’ and ‘Null Distribution2’). The former is a conventional tracking algorithm, whereas the latter reconstructs streamlines by random tracking. The ‘iFOD2’ algorithm (Second-order Integration over FODs) is a probabilistic algorithm which uses a Bayesian approach to account for more than one fiber-orientation within each voxel and takes as input a FOD image represented in the spherical harmonic basis. Candidate streamline paths are drawn based on short curved ‘arcs’ of a circle of fixed length (the step-size), tangent to the current direction of tracking at the current points rather than stepping along straight-line segments (94). The ‘Null Distribution2’ algorithm does not use any image information relating to fiber orientations. This algorithm reconstructs connections based on random orientation samples, identifying voxels where the diffusion data is providing more evidence of connection than that expected from random tracking (95). As this random tracking relies on the same seed and target regions as the ‘iFOD2’ algorithm, we can directly compare the number of reconstructed streamlines between the two tracking algorithms. We used the following parameters for fiber tracking (53,54,96): randomly placed 5000 seeds for each voxel in the ROI, a step length of 0.6 mm, FOD amplitude cutoff of 0.05 and a maximum angle of 45 degrees between successive steps. We applied the anatomically-constrained variation of this algorithm, whereby each participant’s five-tissue-type segmented T1 image provided biologically realistic priors for streamline generation, reducing the likelihood of false positives (86). The set of reconstructed connections were refined by removing streamlines whose length was 3 SD longer than the mean streamline length or whose position was more than 3

SD away from the mean position (97,98). To calculate a streamline’s distance from the core of the tract we resampled each streamline to 100 equidistant nodes and treat the spread of coordinates at each node as a multivariate Gaussian. The tract’s core was calculated as the mean of each fibers x, y, z coordinates at each node.

#### Data analysis

##### Testing the presence of reciprocal connections between ROIs

We independently tested the presence of 1) individually defined hMT+/V5 – hPT connections, 2) group-level hMT+/V5 – hPT connections and 3) group-level hPT– FFA connections. No agreement on statistical thresholding of probabilistic tractography exists, but previous studies have considered a connection reliable at the individual level when it had a minimum of 10 streamlines (53,54,96,99,100). However, setting the same absolute threshold for different connections does not take into account that the probability of connections drops exponentially with the distance between the seed and target regions (101), or the difficulty to separate real from false connections (102). To take these into account, we compared the number of streamlines reconstructed by random tracking (‘Null Distribution2’ algorithm), with those generated by conventional tracking (‘iFOD2’ algorithm) (95,103). Since both algorithms conduct tractography using the same seed and target regions, we can directly compare the number of reconstructed streamlines between them without correcting for a possible difference in the distance between the seed and target regions or their sizes (95).

As done in previous studies (96,100), we calculated the logarithm of the number of streamlines [log(streamlines)] to increase the likelihood of obtaining a normal distribution, which was tested before application of parametric statistics using the Shapiro-Wilk test in RStudio (104). The log-transformed number of streamlines were compared using two-sided paired t-tests. To control for unreliable connections, we calculated the group mean and standard deviation (SD) of the log-transformed number of streamlines for each connection and we discarded participants whose values were more than 3SDs away from the group mean for the respective connection. Connections were only considered reliable when the number of streamlines reconstructed with the ‘iFOD2’ algorithm were higher than the ones obtained with the ‘null distribution’ algorithm. Significance was thresholded at p = 0.05 Bonferroni-corrected for multiple comparisons, p = 0.008 (three connections, two hemispheres).

In case we found no difference between the number of streamlines derived from a null distribution and the number of streamlines generated with the ‘iFOD’ algorithm, these results were additionally tested with Bayesian analyses (e.g., see for instance (105)). Such an analysis was based on the t-values obtained with the t-tests mentioned above and on a Cauchy prior centered on zero (scale = 0.707) representing the null hypothesis. The Bayes factor (BF) values obtained with this analysis represent a measure of how strongly the data supports the null hypothesis of no effect (i.e., low BF values, < 1). The Bayesian analysis was performed using JASP (JASP Team, 2019).

##### Overlap of hMT+/V5 – hPT connections with the IFOF and ILF

To assess whether hMT+/V5 – hPT connections followed the same trajectory as the Inferior Longitudinal Fasciculus (ILF) or the Inferior Frontal Occipital Fasciculus (IFOF), two major white matter bundles that connect the occipital lobe with the temporal and frontal lobes respectively (51,65), we computed the spatial overlap between these bundles. The Dice Similarity Coefficient (DSC) (106) was used as a metric to evaluate the overlap of hMT+/V5 – hPT connections with the ILF and the IFOF, in each participant and hemisphere separately. The DSC measures the spatial overlap between regions A and B, and it is defined as DSC(A,B)= 2(A∩B)/(A+B) where ∩ is the intersection. We calculated the DSC of hMT+/V5 – hPT connections and the ILF, using as region A the binarized tract-density images of hMT+/V5 – hPT connections after transforming them from the template diffusion space to the standard space. Region B was the ILF from the JHU-DTI-based white-matter atlas available in FSL thresholded at a probability of 25%. The same procedure was applied to determine the overlap of hMT+/V5 – hPT connections and the IFOF.

Additionally, individual tract-density images for hMT+/V5 – hPT connections were binarized, summed and thresholded at >9 subjects. The overlap of such hMT+/V5 – hPT maps with the ILF and the IFOF was also computed.

##### Laterality of hMT+/V5 – hPT connections

We assessed the laterality of hMT+/V5 – hPT connections by means of the laterality index (LI). To compute the LI of hMT+/V5 – hPT connections derived from individually defined hMT+/V5 and hPT, we first calculated the connectivity index (CI).

As the number of streamlines connecting two regions strongly depends on the number of voxels in the seed and target masks and we conducted the probabilistic tracking in symmetric mode (from the seed to the target and from the target to the seed), the CI was determined by the number of streamlines from the seed that reached the target divided by the product of the generated sample streamlines in each seed/target voxel (5000) and the number of voxels in the respective seed/target mask (96,100). To increase the likelihood of gaining a normal distribution, log-transformed values were computed (96,100) as follows:

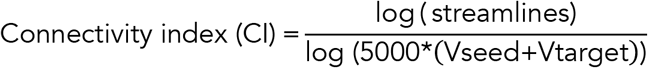

Subjects whose connectivity index was more than 3SDs away from the group mean were considered outliers (96,100).

Once the CI was computed, the laterality index was defined as the subtraction between the connectivity index in the right and left hMT+/V5 – hPT connections, divided by their addition:

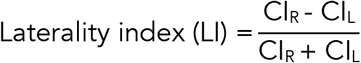

The laterality index varies between −1 and 1, for completely left- and right-lateralized connections, respectively.

##### Testing whether hMT+/V5 – hPT connections are different when relying on individual or group-level hMT+/V5 and hPT

We analyzed the impact that group-level localization of hMT+/V5 and hPT (compared to individually localized regions) had in the reconstruction of hMT+/V5 – hPT connections, as we aimed to 1) investigates the reproducibility of the connection under these two approaches to define the ROIs and 2) assess the existence of hMT+/V5 – hPT connections in Dataset 2 (HCP) where the individual definition of hMT+/V5 and hPT is not possible. We first investigated whether hMT+/V5 – hPT tracts existed when the location of the ROIs was derived from group-averaged functional data, as described in the section “Testing the presence of connections”. Then, the impact was assessed by means of macrostructural and microstructural metrics.

Macrostructural characterization included the comparison of the connectivity index between hMT+/V5 – hPT tracts derived from individual and group functional data, whereas we relied on diffusivity measures derived from fiber orientation distribution for the microstructure assessment.

###### Connectivity index

Connectivity indexes between connections derived from individual and group-level ROIs were compared using two-sided paired t-tests, after testing for normality using the Shapiro–Wilk test. Significance threshold was set at p = 0.025 (right and left hemisphere).

As the connectivity index is highly influenced by the distance between the seed and target regions, we assessed differences in hMT+/V5 – hPT distance between individual or group-level definition of ROIs. Distance in hMT+/V5 – hPT connections was defined as the Euclidian distance between hMT+/V5 and PT coordinates in each subject. Distance values were normally distributed after a log-transformation and we compared them using two-sided paired t-tests. If differences in distance were found, the distance-corrected connectivity index was calculated, replacing the number of streamlines with the product of the distance and the number of streamlines (d*streamlines). Similar approaches have been used by other studies to take into account the effect of distance in the number of streamlines generated between two regions (54,107,108).

###### FOD-derived microstructural metrics

To independently characterize microstructural estimates of streamlines connecting hMT+/V5 and hPT, and isolate the contribution of crossing fibers within the same voxels, we extracted FOD-derived metrics. FOD-derived measures aim at obtaining tract-specific measures, as opposed to voxel-specific tensor-derived metrics (e.g., fractional anisotropy), for which the information of different fiber populations is averaged within a voxel. Because of current limits in DWI spatial resolution, 90% of white matter voxels in the brain include complex fiber configurations (e.g. kissing, fanning, crossing fibers) (109), and the Gaussian assumption of the tensor model is therefore suboptimal to describe tissue microstructure in these voxels (110–113).

The extracted FOD-derived quantitative metrics of microstructural diffusivity were fiber density (FD), fiber-bundle cross-section (FC) and fiber density and cross-section (FDC) following the Fixel-based analysis pipeline (www.mrtrix.org). FD estimates the intra-axonal volume of a fiber pathway within a voxel and was calculated as the integral of the FOD along a particular direction. This metric is proportional to the intra-axonal volume of axons aligned in that direction and is therefore sensitive to alterations at the fixel-level (83). FC reflects changes in a fiber bundle’s intra-axonal volume that are manifested as a difference in the number of voxels/spatial extent that the fiber bundle occupies (cross-sectional area) (90). FCs for each fixel were calculated in the plane perpendicular to the fixel direction and were estimated by using the non-linear warps used to register each subject’s FOD to the template. Lastly, multiplying FD and FC we computed the metric FDC, which combines both sources of information (within-voxel FD and changes in FC). These estimates, were computed in all the voxels that were crossed by fibers connecting hMT+/V5 and hPT and then averaged for each bundle. To compare microstructural diffusivity metrics between tracts derived from subject-versus group-specific ROIs, the same statistical procedure as for the connectivity index was used.

### Dataset 2: HCP

We investigated the reproducibility of hMT+/V5 – hPT connections using an independent dMRI dataset that involves a large number of participants. Minimally processed dMRI data from the new subjects (n=236) in the HCP S1200 release was used (114) (https://www.humanconnectome.org/study/hcp-young-adult/document/1200-subjects-data-release). Participants who did not complete the diffusion imaging protocol (n=51), gave positive drug/alcohol tests (n=19), and had abnormal vision (n=1) were excluded from the analysis. Taking into account the high number of siblings in the HCP sample and the fact that this might spuriously decrease the variance due to the structural and functional similarity, we only selected unrelated participants and kept one member of each family (from the 86 siblings 47 participants were excluded). The selection resulted in 114 healthy participants (50 women; mean age ± SD, 28.6 ± 3.7; range, 22–36).

#### Image acquisition and processing

The minimally processed structural data included T1-weighted high-resolution MPRAGE images (voxel size = 0.7 mm isotropic) corrected for gradient distortions and for low-frequency field inhomogeneities. The diffusion data was constituted by a multi-shell acquisition (b-factor = 1000, 2000, 3000 s/mm2) for a total of 90 directions, at a spatial resolution of 1.25 mm isotropic. Minimal processing included intensity normalization across runs, echo planar imaging (EPI) distortion correction and eddy current/motion correction. Further details on image acquisition can be found elsewhere (https://www.humanconnectome.org/storage/app/media/documentation/s1200/HCP_S1200_Release_Appendix_I.pdf).

The preprocessing of the diffusion images in this dataset is highly similar to the processing of images in Dataset 1. The only difference in the processing of data arises from the multi-shell nature of the acquisition as opposed to the single-shell acquisition of Dataset 1. Given the sample size, we selected a subset of 60 participants (~ half of the total sample) to create a representative population template and white matter mask (103). We used this template to normalize the white matter intensity of all 114 participants’ dMRI volumes.

##### Preparation of ROIs and probabilistic tractography

Individual definition of hMT+/V5 and hPT was not possible, since the HCP scanning protocol does not include any motion localizer experiment. Hence, inclusion regions for tractography were derived from group-level hMT+/V5 and hPT, as described in Dataset 1. The ROIs were transformed to template diffusion space, where we conducted tractography using the same procedure used in Dataset 1.

#### Data analysis: Testing the presence hMT+/V5 – hPT connections

We assessed the existence of reciprocal connections between group-level hMT+/V5 and hPT as described in Dataset 1.

## ACKNOWLEDGEMENTS

The project was funded by the ERC starting grant MADVIS (Project: 337573), the Belgian Excellence of Science program from the FWO and FRS-FNRS (Project: 30991544) and a “mandat d’impulsion scientifique (MIS)” from the FRS-FNRS awarded to OC. Computational resources have been provided by the supercomputing facilities of the Université catholique de Louvain (CISM/UCL) and the Consortium des Équipements de Calcul Intensif en Fédération Wallonie Bruxelles (CÉCI) funded by the Fond de la Recherche Scientifique de Belgique (F.R.S.-FNRS) under convention 2.5020.11 and by the Walloon Region. AG is supported by the Wallonie Bruxelles International Excellence Fellowship and the FSR Incoming PostDoc Fellowship by Université Catholique de Louvain. OC is a research associate and MR a doctoral fellow supported by the Fond National de la Recherche Scientifique de Belgique (FRS-FNRS).

